# Brain activation and connectivity in anorexia nervosa and body dysmorphic disorder when viewing bodies: relationships to clinical symptoms and perception of appearance

**DOI:** 10.1101/2020.02.12.934083

**Authors:** Teena D Moody, Francesca Morfini, Gigi Cheng, Courtney L Sheen, Wesley Kerr, Michael Strober, Jamie D Feusner

**Author notes:** **Address for correspondence:** Teena D Moody, https://orcid.org/0000-0001-7067-9512?lang=en; UCLA Semel Institute 27-465, Los Angeles, CA 90095. **Disclosures:** The authors report no competing interests. The funding sources had no participation in conducting the study. The UCLA Institutional Review Board (IRB) approved the study. **Funding:** National Institutes of Health, USA grant R01MH093535 (Feusner) & T32 GM08042 (Porterra).

## Abstract

Anorexia nervosa (AN) and body dysmorphic disorder (BDD) are characterized by distorted perception of appearance, yet no studies have directly compared the neurobiology associated with body perception. We compared brain activation and connectivity in relevant networks when viewing images of others’ bodies and tested their relationships with clinical symptoms and subjective appearance evaluations. We acquired fMRI data from 64 unmedicated females (20 weight-restored AN, 23 BDD, 21 controls) during a matching task using photos of others’ bodies that were unaltered or spatial-frequency filtered. With general linear model and independent components analyses we compared brain activation and connectivity in visual, striatal, and parietal networks and performed univariate and partial least squares multivariate analyses to investigate relationships with clinical symptoms and appearance evaluations. BDD but not AN demonstrated hypoactivity in dorsal visual and parietal networks compared to controls. Yet, AN and BDD showed partially overlapping patterns of hyperconnectivity in the dorsal visual network and hypoconnectivity in parietal network compared with controls. Further, there were significant activity and connectivity differences between AN and BDD in both networks. In both groups, activity and/or connectivity were associated with symptom severity and appearance ratings of others’ bodies. AN and BDD demonstrate both distinct and partially overlapping aberrant neural phenotypes involved in body processing and visually encoding global features. Nevertheless, in each disorder, aberrant activity and connectivity show relationships to clinically relevant symptoms and subjective perception. Results have implications for understanding distinct and shared pathophysiology underlying perceptual distortions for appearance and may inform future novel treatment strategies.

## INTRODUCTION

Anorexia nervosa (AN) and body dysmorphic disorder (BDD) are considered to be different nosological entities but they share a distorted perception of appearance (Jolanta and Rabe-Jablonska Jolanta 2000), obsessive and compulsive symptoms (S Ruffolo et al. 2006; Kittler, Menard, and Phillips 2007; Madsen, Bohon, and Feusner 2013), and poor insight (Grant, Kim, and Eckert 2002; Deckersbach et al. 2000). BDD is characterized by anomalous perception of features of the face, head, or body parts (Ines Kollei et al. 2012), in addition to preoccupation with these appearance concerns and repetitive behaviors such as checking or attempts to fix or hide appearance features (American Psychiatric Association 2013). AN is defined by a low body mass, an indifference to the consequences of low body mass and frequently an insistence that further weight loss is needed. In AN, the appearance concerns are more often about areas of the body typically linked to weight, such as stomach and waist, whereas in BDD the concerns are more frequently about facial features, hair, and skin (Toh et al. 2019). Individuals with BDD tend to have lower insight and are more likely to have delusional beliefs about their appearance than those with AN (Andrea S. Hartmann et al. 2013). However, BDD and AN generally share reduced appearance satisfaction, frequent appearance evaluations, attitudes towards one’s self, more body areas of dissatisfaction, and greater preoccupation with being overweight, compared with controls, as well as high bi-direction comorbidity (A. S. Hartmann et al. 2015; Hrabosky et al. 2009; I. Kollei et al. 2017). Among persons with AN, 25–39% are diagnosed with lifetime BDD and 32% of those with BDD have a lifetime eating disorder (Grant, Kim, and Eckert 2002; Jolanta and Rabe-Jablonska Jolanta 2000; S Ruffolo et al. 2006). Moreover, 30% of individuals with BDD have significant weight-related appearance concerns (Kittler, et al., 2007). Thus, AN and BDD may have partially overlapping features including body-image aberrations and could share genetic, epigenetic, or other susceptibilities. Hence, we pose this question: Do AN and BDD share neural phenotypes that might account for the overlap in clinical phenomenology and cross-disorder comorbidity?

Lines of evidence suggest that AN and BDD may share visual and visuospatial experiences (Madsen, Bohon, and Feusner 2013); specifically, that details of physical appearance are perceived as abnormally prominent, rather than considered in a global context. Neuropsychological studies of the two disorders suggest imbalances in global versus local visual processing (Deckersbach et al. 2000; Lang et al. 2016; Sherman et al. 2006).

Previous investigations of neural visual system functioning in AN and BDD provide evidence of abnormalities that could contribute to these perceptual distortions (Beilharz et al. 2017; J. Feusner, Deshpande, and Strober 2017; Pietrini et al. 2011; Groves, Kennett, and Gillmeister 2019). In a combined analysis of functional magnetic resonance imaging (fMRI) and EEG data (Li, Lai, Bohon, et al. 2015), persons with AN and BDD demonstrated hypoactivity in the dorsal visual stream for face stimuli and in the dorsal visual stream and occipital fusiform cortex when viewing house stimuli. These differences were apparent for low spatial frequency (LSF) images of faces and houses. This hypoactivity in dorsal visual stream and early visual areas might result in deficiencies in the holistic appraisal of visual stimuli, in turn, interfering with formation of a perceptual whole. Yet, an fMRI connectivity study of face processing in AN and BDD identified *different* patterns in lower-level face processing systems (occipital and fusiform face areas) although there was *similar* higher-order connectivity between fusiform face area and precuneus/posterior cingulate (Moody et al. 2015). Thus, the few direct comparisons of neural phenotypes in AN and BDD suggest partial overlap, with both distinct and similar visual system neural patterns for LSF appearance (face) and non-appearance-related (house) stimuli.

Whether overlapping and distinct patterns in AN and BDD exist when viewing symptom-relevant stimuli of *bodies* has yet to be explored. In AN, fMRI studies of *own-body* processing have found decreased activity in inferior parietal lobule and precuneus (Mohr et al. 2010; Sachdev et al. 2008; Vocks, Busch, Schulte, et al. 2010; Vocks, Busch, Gronemeyer, et al. 2010), although one study found increased inferior parietal lobule activity (Wagner et al. 2003). FMRI studies in AN using *other-body* stimuli have been less consistent. Weaker activation to other-body images was found in occipitotemporal cortex (including extrastriate body area) and superior parietal lobule (Uher et al. 2005). Another study found increased activation of inferior and superior parietal lobule, inferior and middle frontal gyri, parahippocampus, and amygdala (Vocks, Busch, Gronemeyer, et al. 2010). These studies suggest abnormalities in body-processing and limbic regions in AN.

Connectivity studies using *other-body* stimuli have found that individuals with AN who have weaker connectivity strength from left fusiform body area to left extrastriate body area demonstrated increased body-size misjudgment (Suchan et al. 2013), whereas another study showed that individuals with AN showed stronger connectivity between precuneus and mid-temporal regions when viewing others’ bodies (Via et al. 2018). These connectivity differences in other-body processing may reflect perceptual abnormalities and contribute to a key symptom of body-size misjudgment. Studies of neural correlates of body processing in BDD have not been conducted, and no study has compared neural correlates of body processing in AN and BDD directly in the same task.

With the aim of elucidating the neural underpinnings of anomalies of perception, the objective of this study was to characterize brain activation and connectivity patterns in AN and BDD compared with healthy controls when viewing *others’ bodies*. We chose *others’* instead of *own* bodies to assess general abnormalities in brain networks responsible for body processing (Li, Lai, Bohon, et al. 2015; Li, Lai, Loo, et al. 2015; Moody et al. 2015) and to minimize the confound of emotional arousal of viewing participants’ own bodies. While bodies are more salient and would be expected to evoke greater emotional arousal in AN than BDD, viewing others’ bodies rather than own bodies was designed to constrain this effect. Importantly, using the same stimuli for all participants also allows for a direct comparison of visual processing in the two diagnostic groups.

We focused on 3 networks: the dorsal visual network, a parietal network including the inferior parietal lobule, and the striatal network. The dorsal visual network includes regions in dorsal visual stream involved in global/configural image processing (Goodale and Milner 1992; Ungerleider and Haxby 1994) that receive magnocellular input conveying LSF information (Merigan and Maunsell 1993). Visual processing abnormalities for BDD and AN previously have been found in dorsal visual stream regions (J. D. Feusner et al. 2007; Li, Lai, Bohon, et al. 2015; Wagner et al. 2003). The dorsal visual network covers the extrastriate body area and portions of precuneus. The parietal network, including inferior parietal lobule and associated regions, is involved in body perception (Engelen et al. 2015; Hodzic et al. 2009), and abnormal functioning has been implicated both in AN (Vocks, Busch, Gronemeyer, et al. 2010; Wagner et al. 2003) and in BDD for faces (J. D. Feusner et al. 2007). Support for testing striatal network involvement comes from studies of face processing in BDD (J. D. Feusner et al. 2010), and of body perception and reward in AN (Bischoff-Grethe et al. 2013; Fladung et al. 2010; Decker, Figner, and Steinglass 2015; McFadden et al. 2014; Frank et al. 2012). Body visual processing is not restricted to these networks; however, given our sample size, we focused on networks previously implicated in AN and BDD.

We hypothesized that AN and BDD would have abnormally low activation and connectivity in the dorsal visual network for LSF images. We predicted greater activation and connectivity in striatal network for normal spatial frequency (NSF) images for both groups compared with controls, as evidenced by previous studies in AN and BDD across different tasks. For the parietal network, we predicted aberrant activity and connectivity in both groups for NSF images; as there was less evidence from previous studies, we did not have directional hypotheses. We also examined associations between symptom severity and network connectivity and activation strength. For LSF, we hypothesized that low insight scores and symptom severity would be associated with lower dorsal visual network activity and connectivity. Post-hoc we investigated multivariate relationships between regions of abnormal activation and connectivity and clinical variables, using partial least squares analyses. As an exploratory analysis, we also examined activation and connectivity for high spatial frequency (HSF) stimuli.

## METHODS

### Study design and participants

Participants included 64 females ages 13-33: 20 weight-restored AN (to avoid confounds of starvation state), 23 BDD, and 21 healthy controls. All were unmedicated, right-handed, recruited from the community, and were matched for age and years of education. All underwent the Mini International Neuropsychiatric Interview (MINI) (Sheehan et al. 1998) and the BDD Diagnostic Module (Phillips, Atala, and Pope 1995). Licensed clinicians (JF and MS) established diagnoses. Illness duration was assessed by the clinician, who determined at what age the participant first had clinically significant symptoms that resulted in impairment of function or significant distress. AN participants had to have previously met full DSM-IV criteria, and currently meet all criteria except for weight and amenorrhea. Psychometrics included the Hamilton Anxiety Rating Scale (HAMA) (Hamilton 2016), Montgomery–Asberg Depression Rating Scale (MADRS) (Montgomery and Asberg 1979), Brown Assessment of Beliefs Scale (BABS) (Eisen et al. 1998), the BDD version of the Yale-Brown Obsessive-Compulsive Scale (BDD-YBOCS) (Phillips et al. 1997), and the Eating Disorder Examination (EDE) Edition 16.0D (Fairburn, Cooper, and M. E. 2008). Subjective ratings of body images with respect to attractiveness, aversiveness, and over/underweight judgements were acquired outside the scanner. The University of California, Los Angeles Institutional Review Board approved the protocol and all participants provided written informed consent. (See Supplemental Materials for more details of ratings and for inclusion/exclusion criteria.)

### Bodies Task

In the scanner, participants performed a task of matching photos of other people’s bodies and, as a control task, matching shapes. The photos were black/white photos of other female and male bodies wearing underwear, with heads cropped, ranging from normal to overweight. The photos were unaltered (normal spatial frequency -NSF) or spatial frequency-filtered to include only LSF and HSF as previously described (Iidaka et al. 2004). The forced-choice, two-sample task consisted of a target body and two selection bodies that appeared simultaneously on the screen for 4 s (Fig. 1). Participants pressed the right or left button to choose the selection body that was identical to the target body. A fixation cross appeared during the inter-stimulus interval of 0.5 s. A total of 48 sets of body stimuli were presented in alternating blocks of NSF, LSF, or HSF, or control, with four sets of images per block. The order of stimuli was counterbalanced across participants using a Latin-squares design.

**Fig. 1.**
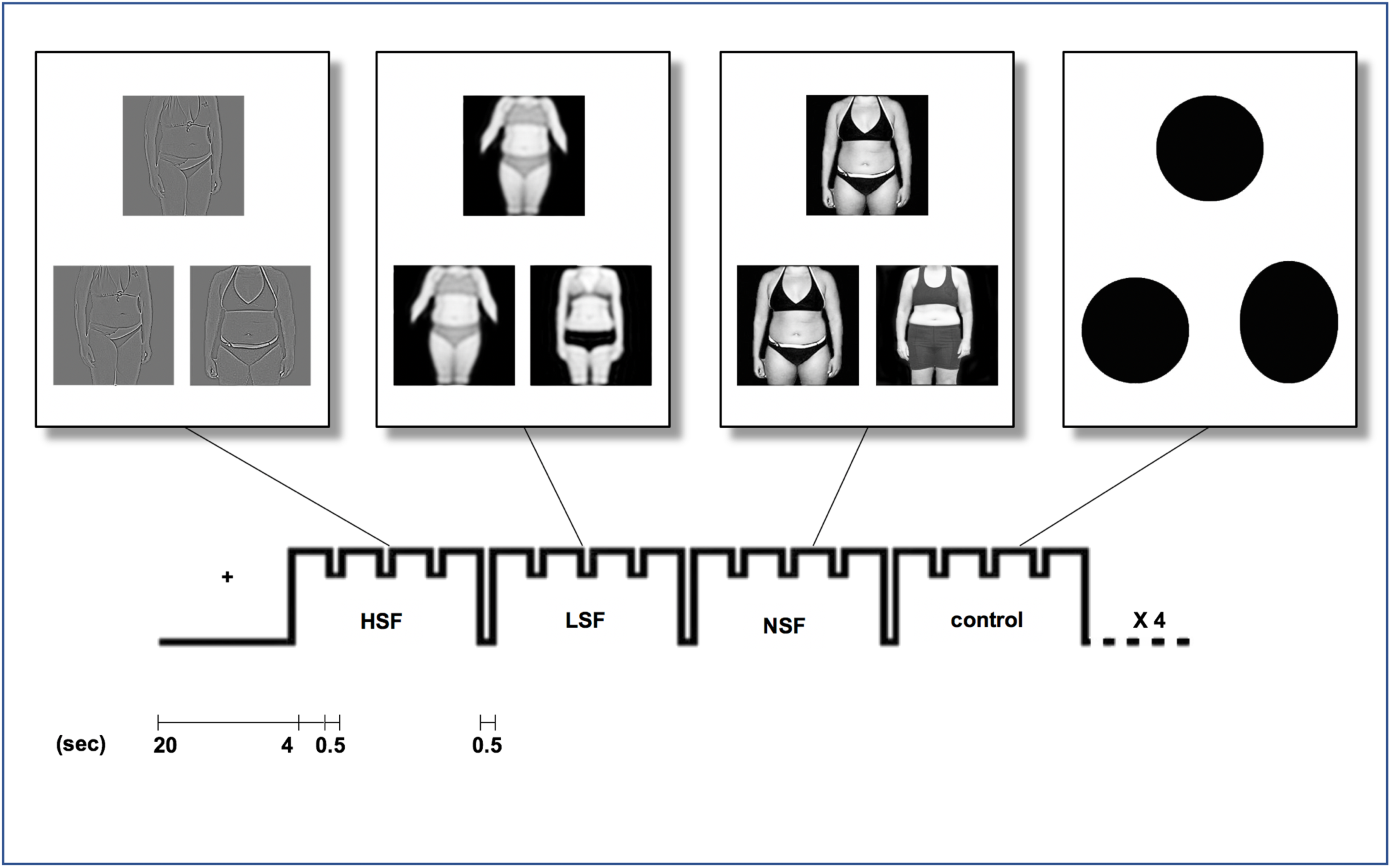
Bodies matching task design. Participants engaged in a task of matching photos of bodies or shapes in the MR scanner. The photos were black and white photos of other people’s bodies (female and male subjects) wearing underwear, ranging from normal to overweight (Moody *et al*., 2017).

### MRI data acquisition and processing

We acquired whole-brain blood oxygenation level dependent (BOLD) fMRI and structural MRI data with on a Siemens TRIO 3-T scanner using a 12-channel coil (described previously (Li, Lai, Bohon, et al. 2015; Moody et al. 2015). We collected functional echo-planar images (EPI) using: repetition time (TR) 2.5 s; echo time (TE) 25 ms; flip angle, 80°; voxel dimensions, 3 × 3 × 3 mm; 0.75 mm gap; field-of-view, 192 mm; matrix, 64 × 64; 133 measurements; 32 slices. Data collected during the first three TRs were discarded for T1 equilibration. The data for each participant therefore contained 30 time points for images of each spatial frequency (NSF, LSF, HSF) and control images. We obtained an MPRAGE T1-weighted image to provide detailed brain anatomy with: TR 1.9 s, TE 2.26 ms, and voxel dimensions 1 × 1 × 1 mm. MRI data were preprocessed using FMRI Expert Analysis Tool (FEAT) version 6.0, FSL (FMRIB’s Software Library; http://www.fmrib.ox.ac.uk/fsl), (see Supplemental Materials). Image preprocessing for connectivity denoised each individual participant’s data using FSL’s AROMA (Pruim et al. 2015).

### Activation

We used general linear models to assess activation patterns among groups and extracted eigenvalues. Contrasts of interest at the subject level were LSF bodies and NSF bodies vs. shapes. We did not have specific hypotheses for HSF analyses, therefore these were exploratory. Group analyses with between- and among-group contrasts, used FMRIB’s Local Analysis of Mixed Effects (FLAME 1+2) to assess differences in activation for LSF, NSF, and HSF body photos. Due to the age range that spanned adolescents and young adults, we used age and age-squared as regressors of non-interest to account for potential linear and nonlinear effects of age. We did not control for body mass index (BMI) because controlling for BMI would remove group effects of interest. (It is important to note that while mean BMI is slightly lower in our AN participants, imaging studies have shown that weight-restored AN recover from brain size differences that have been found in those who are not weight-restored (King et al. 2015; Seitz et al. 2014).) Z-statistic images were cluster-thresholded at z>2.3 with a corrected cluster significance threshold of p<0.05; where multiple tests were performed, multiple-correction adjustments (Bonferroni correction) were applied. Network masks derived from the NSF, LSF, and HSF connectivity analyses identified the dorsal visual, parietal, and striatal networks of interest for activation analyses, following the method described by (Filippini et al. 2009). (Network mask details are in Supplemental Materials).

### Connectivity

We used independent components analysis (ICA) to assess temporally-related neuronal activation patterns in brain networks, henceforth referred to as *functional connectivity*. We used Multivariate Exploratory Linear Optimized Decomposition into Independent Components (MELODIC) (Filippini et al. 2009; Jenkinson et al. 2012; S. M. Smith et al. 2004) and dual regressions to perform voxel-wise comparisons of functional connectivity (Supplemental Materials). We used nonparametric permutation testing, FSL’s randomise, (Stephen M. Smith and Nichols 2009) to determine statistically significant differences among groups, controlling for age and age-squared, and applying family-wise error rate correction of p<0.05.

### Associations with Symptom Severity

We tested hypotheses regarding the dorsal visual network for LSF activation and connectivity analyses, using psychometric instruments that have been validated in each respective diagnostic group as covariates: BABS and BDD-YBOCS scores were used for BDD, and BABS and EDE for AN. BABS is a metric of insight, BDD-YBOCS measures BDD symptom severity, and EDE measures eating disorder severity. We controlled for age and age-squared.

### Multivariate association of psychometrics scores and appearance ratings

As post hoc analyses, we used partial least squares (PLS) regression to explore associations between eigenvalues (activation) or coherence values (functional connectivity) and symptom severity scores or appearance ratings of others’ bodies. Eigenvalues and connectivity values were tested separately and were extracted from multiple clusters found to be different in between-group analyses. BABS and EDE shape-concern scores were used for AN, and BABS and BDD-YBOCS scores were used for BDD. Subjective ratings of the appearance of the body photos used in the fMRI task were obtained outside the scanner, after MRI data collection, for attractiveness, aversiveness, and over/underweight judgments as described in Supplemental Materials. These exploratory, post hoc analyses were not corrected for multiple comparisons.

## RESULTS

### Participants

BDD, AN, and control groups did not differ in mean age or years of education (Table 1). ANOVA/t-tests revealed significant differences among groups for BMI [F (2,61)=4.66, p<0.013], HAMA [F (2,61)=11.23, p<0.0001], and BABS [t (19)=-2.82, p=0.011].

**Table 1.** Demographics and Psychometrics. All participants were unmedicated and all AN were weight-restored. Results are presented as mean values ± standard deviations. * Significant at p<0.05. ^The value of BABS was missing for three participants with AN. Abbreviations: BMI, body mass index; HAM-A, Hamilton Anxiety Rating Scale; BABS, Brown Assessment of Beliefs Scale; YBC-EDS, Yale-Brown-Cornell Eating Disorder Scale; EDE, Eating Disorder Examination; BDD-YBOCS, Body Dysmorphic Disorder Yale Brown Obsessive Compulsive Scale; LSF, low spatial frequency; NSF, normal spatial frequency.

| Demographics | AN | BDD | CON | Stats<br>(F or T) | P-value |
| --- | --- | --- | --- | --- | --- |
| **Females** | **20** | **23** | **21** |  |  |
| **Age, years** | 23.3±3.3 | 23.8±5.0 | 21.9±4.7 | 1.07 | 0.35 |
| **Age range (min,max)** | (18,30) | (18,33) | (13,33) |  |  |
| **Years of education (n)** | 15.3±2.3(19) | 15.6±3.4(22) | 14.2±2.6 | 1.58 | 0.21 |
| **Illness duration, months** | 91.3±59.1(19) | 130.6±79.3(22) |  | -1.82 | 0.08 |
| **BMI** | 20.4±1.4 | 22.5±2.9 | 22.7±3.3 | 4.66 | 0.013* |
| **Lowest lifetime BMI** | 16.1±1.6(16) |  |  |  |  |
| **Subtype AN (n)** | Restricting (16)<br>Binge/purge (4) |  |  |  |  |
| **Type of concern BDD (n)** |  | Face only (8)<br>Body only (0)<br>Body and Face (15) |  |  |  |
| **Co-morbidities** |  |  |  |  |  |
| Major depressive disorder | 1 | 6 |  |  |  |
| Dysthymia | 2 | 3 |  |  |  |
| Generalized anxiety disorder | 4 | 4 |  |  |  |
| Social anxiety disorder |  | 4 |  |  |  |
| Panic disorder |  | 2 |  |  |  |
| Agoraphobia |  | 3 |  |  |  |
| None | 13 | 10 |  |  |  |
| **Symptoms severity self-reports** |  |  |  |  |  |
| BABS | 10.2±7.2 | 15.3±2.6 |  | -2.82 | 0.011* |
| BDD-YBOCS |  | 30.6±6.6 |  |  |  |
| EDE total scores | 2.9±0.5 |  |  |  |  |
| Shape-concern scores | 3.7±1.7 |  |  |  |  |
| HAM-A | 7.6±6.4 | 10±7.5 | 1.9±1.3 | 11.23 | <0.001* |
| MADRS | 10.6±9.1 | 16±8.5 | 1.1±1.8 | 23.39 | <0.001* |
| **Ratings** | **AN (n=15)** | **BDD (n=18)** | **CON (n=19)** |  |  |
| LSF - Aversion | 3.4±2.0 | 3.9±2.4 | 1.9±1.3 | 5.46 | 0.01* |
| - Attractiveness | 4.6±1.3 | 4.6±1.3 | 5.0±1.5 | 0.39 | 0.68 |
| - Weight perception | 2.9±1.6 | 2.6±1.5 | 1.3±0.9 | 7.14 | 0.002* |
| NSF - Aversion | 3.5±1.8 | 3.5±2.0 | 2.2±1.2 | 3.48 | 0.04* |
| - Attractiveness | 5.0±1.4 | 5.0 ±0.9 | 4.7±1.4 | 0.33 | 0.72 |
| - Weight perception | 2.8±1.4 | 2.5 ± 1.0 | 1.7±1.1 | 4.83 | 0.012* |

### Behavioral

Neither mean accuracy (95.7±3.1, p=0.52) nor response time (1.2±0.26 sec, p=0.91) differed significantly among groups.

### Activation

#### Dorsal Visual Network–LSF

(Fig. 2a, Table 2a) BDD demonstrated lower activation than control in bilateral superior lateral occipital cortex, bilateral precuneus, and left superior parietal lobule. There were no significant differences between AN and controls. BDD demonstrated greater activation than AN in right lingual gyrus.

**TABLE 2a.** Coordinates of brain activation differences at LSF.

| **LSF** | **Z max** | **x** | **y** | **z** | **Regions** |
| --- | --- | --- | --- | --- | --- |
| AN ≠ BDD ≠ CON (F-test) |  |  |  |  |  |
|  | 10.00 | 22 | -92 | 0 | R OP, R LOC |
|  | 5.26 | 50 | 40 | 12 | R FP, R IFG |
|  | 6.00 | 18 | -32 | -2 | R Th, R HP |
|  | 4.96 | 26 | -46 | 14 | R PCC, R LING |
|  | 4.08 | 50 | -20 | 40 | R PCG |
|  | 5.15 | -18 | -32 | 0 | L Th |
| BDD<CON (t-test) |  |  |  |  |  |
|  | 5.06 | -24 | -60 | 64 | L LOC, L SPL |
|  | 4.77 | -4 | -74 | 40 | L PCUN |
|  | 4.04 | -30 | -78 | 38 | L LOC |
|  | 4.44 | 8 | -72 | 44 | R PCUN, R LOC |
|  | 3.97 | 10 | -44 | 72 | R PoCG |
| BDD>AN (t-test) |  |  |  |  |  |
|  | 3.80 | 12 | -74 | -10 | R LING |

**Fig. 2.**
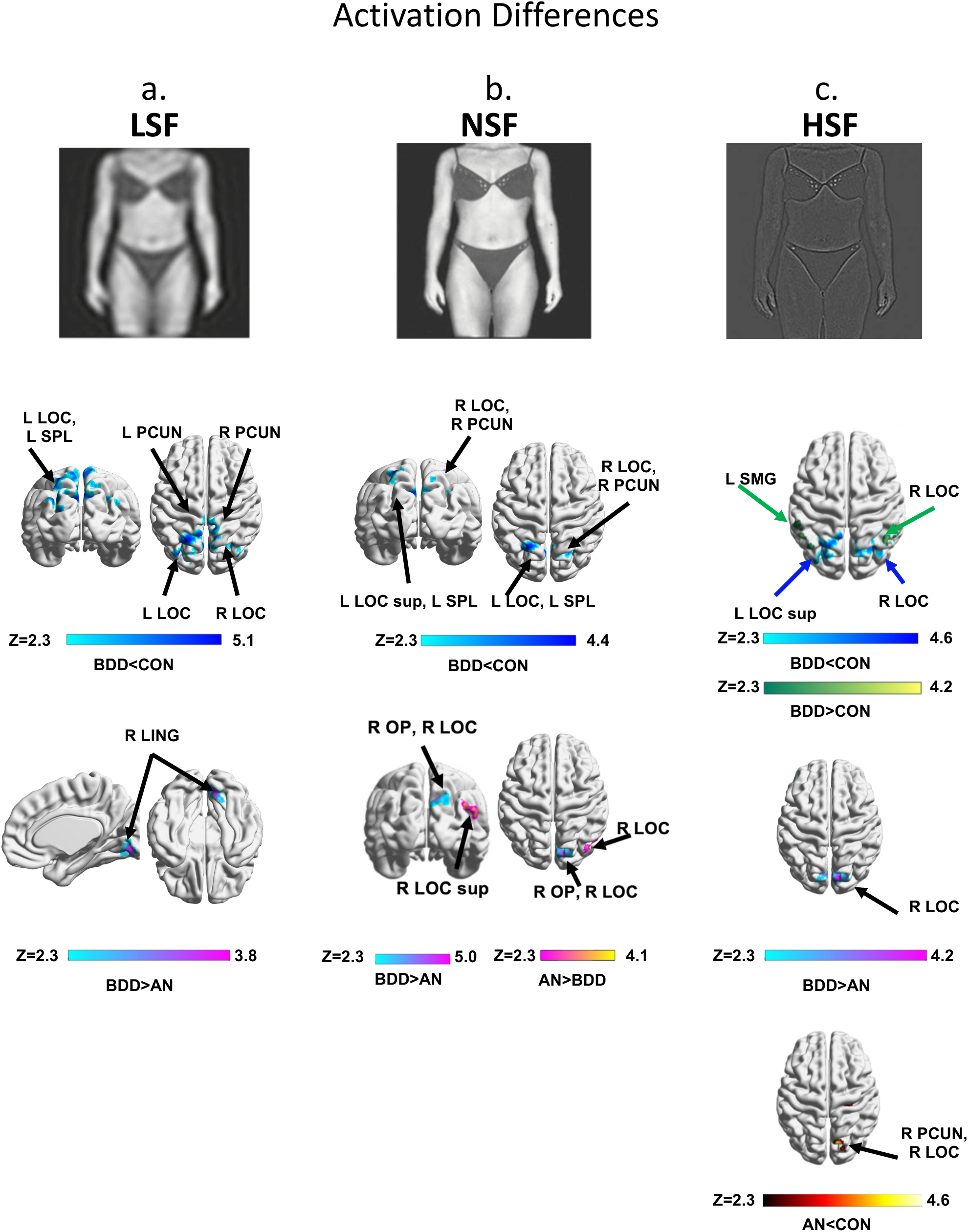
BOLD activity differences with stimuli. ANOVA identified regions differing among the three groups, Z>2.3, using threshold-free cluster enhancement with a family-wise error correction at p<0.05, followed by pairwise group comparisons within significant regions from the ANOVA. (2a) Regions of differences in activation within the dorsal visual network for low spatial frequency (LSF), (2b) within the parietal network for normal spatial frequency (NSF) images, and (2c) regions for high spatial frequency (HSF) images. Examples of bodies stimuli used in the experiment are shown for (2a) LSF, (2b) NSF, and (2c) HSF. Abbreviations: AN, anorexia nervosa; BDD, body dysmorphic disorder; CON, controls; F, frontal; L, left; P, posterior; R, right; V, ventral; Z, z scores. All brain figures were created with BrainNet Viewer (Xia, Wang, and He 2013).

#### Striatal Network-NSF

There were no significant differences between BDD, AN, and control.

#### Parietal Network–NSF

(Fig. 2b, Table 2b) There was lower activation in BDD than controls in bilateral superior lateral occipital cortex, right precuneus, and left superior parietal lobule. There were no significant differences between AN and control. BDD showed lower activation than AN in right superior lateral occipital cortex and greater activation in right occipital pole and superior lateral occipital cortex.

**TABLE 2b.** Coordinates of brain activation differences at NSF.

| **b. NSF** | **Z max** | **x** | **y** | **z** | **Regions** |
| --- | --- | --- | --- | --- | --- |
| AN ≠ BDD ≠ CON (F-test) |  |  |  |  |  |
|  | 9.86 | 22 | -92 | 0 | R OP, R LOC |
|  | 5.54 | 42 | 12 | 28 | R IFG |
|  | 4.99 | -54 | 36 | 14 | L FP, L IFG |
|  | 4.57 | 2 | 58 | -16 | R FP |
|  | 5.29 | -22 | -2 | -16 | L AMYG, L PHC |
|  | 9.86 | 22 | -92 | 0 | R OP, R LOC |
| BDD<CON (t-test) |  |  |  |  |  |
|  | 3.92 | 20 | -68 | 42 | R LOC sup R PCUN |
|  | 4.41 | -26 | -60 | 64 | L LOC sup, L SPL |
| BDD>AN (t-test) |  |  |  |  |  |
|  | 4.97 | 12 | -88 | 40 | R OP, R LOC sup |
| BDD<AN (t-test) |  |  |  |  |  |
|  | 4.15 | 50 | -78 | 28 | R LOC sup |

### Exploratory HSF Activation (Fig2c, Table 2c)

#### Dorsal Visual Network

BDD showed lower activation than controls bilaterally in superior lateral occipital cortex and higher activation than control in right lateral occipital cortex and left supramarginal gyrus. AN showed lower activation than controls in right precuneus, right lateral occipital cortex, and right pre- and postcentral gyri. There was higher activation for BDD than AN in right cuneus, bilaterally in lateral occipital cortex, right lingual gyrus, and left inferior frontal gyrus.

**TABLE 2c.** Coordinates of brain activation differences at HSF.

| c. HSF | Z max | x | y | z | Regions |
| --- | --- | --- | --- | --- | --- |
| AN ≠ BDD ≠ CON (F-test) |  |  |  |  |  |
|  | 10.00 | 38 | -66 | -10 | R OF R LOC |
|  | 6.66 | 42 | 12 | 30 | R MFG, R IFG |
|  | 5.04 | -50 | 32 | 30 | L MFG, L FP |
|  | 4.98 | 2 | 56 | 0 | R FP, R PCG |
|  | 3.93 | -24 | -46 | 66 | L SPL |
|  | 5.22 | 2 | 20 | 54 | R SFG |
|  | 4.16 | 24 | 42 | -16 | R FP |
| BDD < CON (t-test) |  |  |  |  |  |
|  | 4.65 | 28 | -60 | 38 | R LOC sup |
|  | 4.25 | -28 | -62 | 54 | L LOC sup |
|  | 3.74 | -30 | -76 | 36 | L LOC sup |
| AN < CON (t-test) |  |  |  |  |  |
|  | 4.57 | 16 | -68 | 40 | R PCUN, R LOC |
|  | 3.94 | 28 | -24 | 56 | R PrCG, R PoCG |
| BDD > AN (t-test) |  |  |  |  |  |
|  | 4.17 | 10 | -82 | 38 | R CUN, R LOC |
|  | 3.69 | -54 | 24 | -2 | L IFG |
|  | 3.81 | 4 | -72 | -6 | R LING<br>L LOC |
| BDD > CON (t-test) |  |  |  |  |  |
|  | 4.15 | 58 | -60 | 38 | R LOC |
|  | 3.41 | -58 | -42 | 56 | L SMG |

The table shows the maximum z-score, x-y-z coordinates, and brain regions of the peak clusters. Results were thresholded at $z > 2.3$ , $p < 0.05$ . Pairwise comparison t-tests are related to testing our specific hypotheses, but are limited to regions within the significant F-test results.
Abbreviations: L = left; R = right; CG = cingulate gyrus; CUN = cuneal cortex; FOC = frontal orbital cortex; FP = frontal pole; HP = hippocampus; IMG = inferior frontal gyrus; LING = lingual gyrus; LOC = lateral occipital cortex; MTG = middle temporal gyrus; MFG = middle frontal gyrus; OP = occipital pole; OF = occipital fusiform; PCC = posterior cingulate; PCG = paracingulate gyrus; PCUN = precuneus; PHC = parahippocampal cortex; PrCG = precentral gyrus; PoCG = postcentral gyrus; SMG = supramarginal gyrus; SPL = superior parietal lobule; Th = thalamus; an = anterior division; po = posterior division; pt = pars triangularis; sup = superior.

#### Striatal and Parietal Networks

BDD showed lower activation than control bilaterally in superior lateral occipital cortex. (Note that for LSF, NSF, and HSF, there were regions within the 3 ICA-derived network masks that overlapped with each other in lateral occipital cortex.)

### Associations between symptom severity and brain activation

#### Dorsal Visual Network–LSF

In BDD, lower activation in right inferior and middle frontal gyri was associated with better insight (lower BABS scores, p=0.006, Fig. 3a). In AN, lower activation in right lingual gyrus and right intracalcarine cortex was associated with worse insight (higher BABS scores) (p=0.024, Fig. 3b). In individuals with AN, worse eating disorder symptoms (EDE total scores) were associated with lower activation in midline intra/supra calcarine cortex and occipital pole (p=6.9×10^−5^, Fig. 3c). A follow-up exploratory analysis found that worse shape concerns (EDE shape-concern subscale scores) were even more strongly associated with lower activity in similar locations and extending to left lingual gyrus (p=4.0×10^−10^, Fig. S3a).

**Fig. 3.**
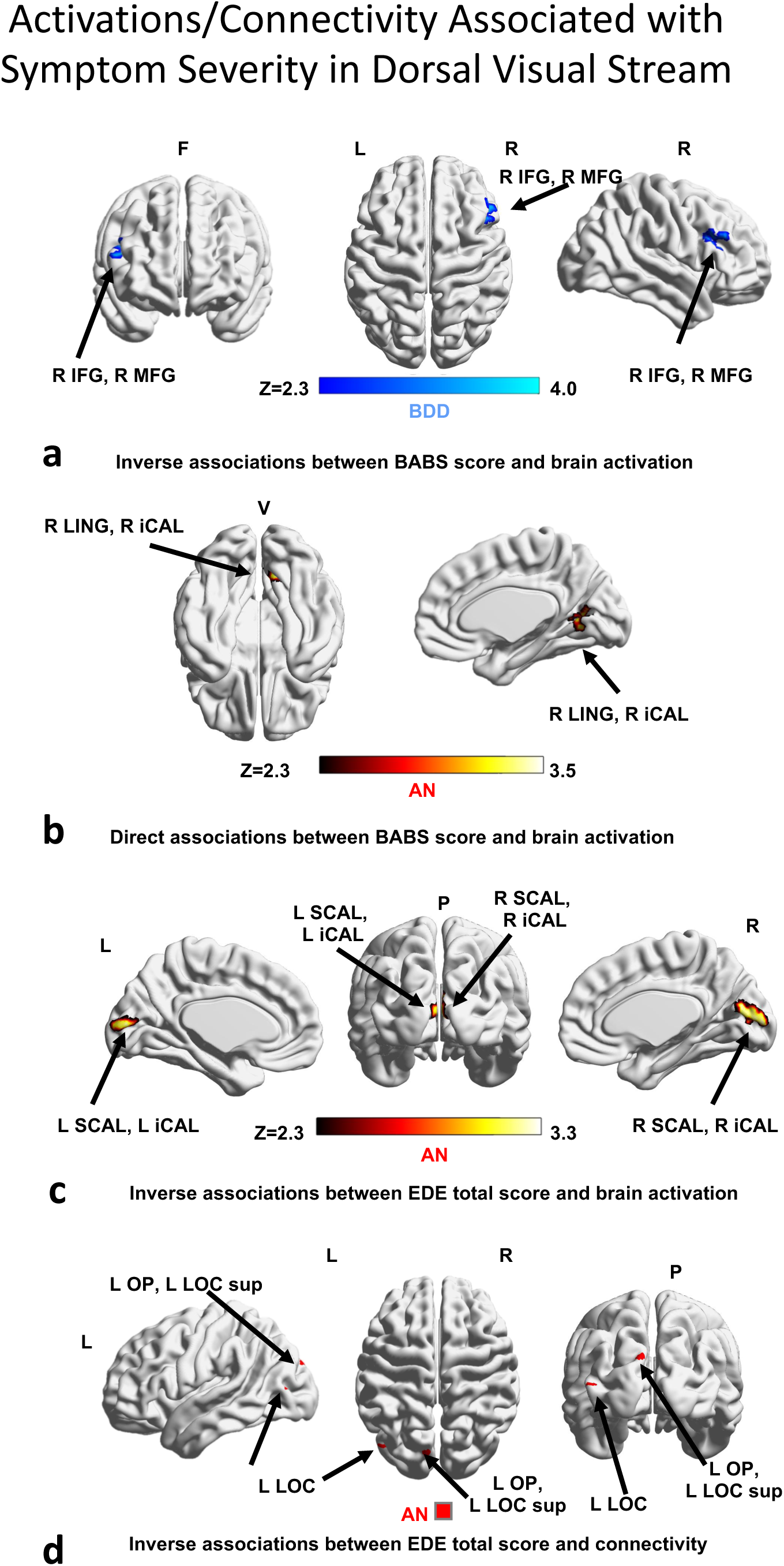
Associations of regions that covary with poor insight and symptom severity scores in dorsal visual stream: (3a) activation inversely associated with BABS scores in BDD; (3b) activation directly associated with BABS scores in AN; (3c) activation inversely associated with EDE score in AN; (3d) connectivity inversely associated with EDE score in AN. Abbreviations: AN, anorexia nervosa; BABS, Brown Assessment of Beliefs Scale; BDD, body dysmorphic disorder; EDE, Eating Disorder Examination; F, frontal; L, left; P, posterior; R, right; V, ventral; Z, z scores.

### Multivariate PLS associations of symptom severity from brain activation

#### Dorsal Visual Network–LSF (Fig. 4a)

In AN, aggregate activation strength was significantly associated with symptom severity (BABS and EDE shape-concern, R^2^= 0.24, p=0.030). In BDD, activation strength showed a trend for association with symptom severity(BABS and BDD-YBOCS) (R^2^=0.17, p=0.054).

**Fig. 4.**
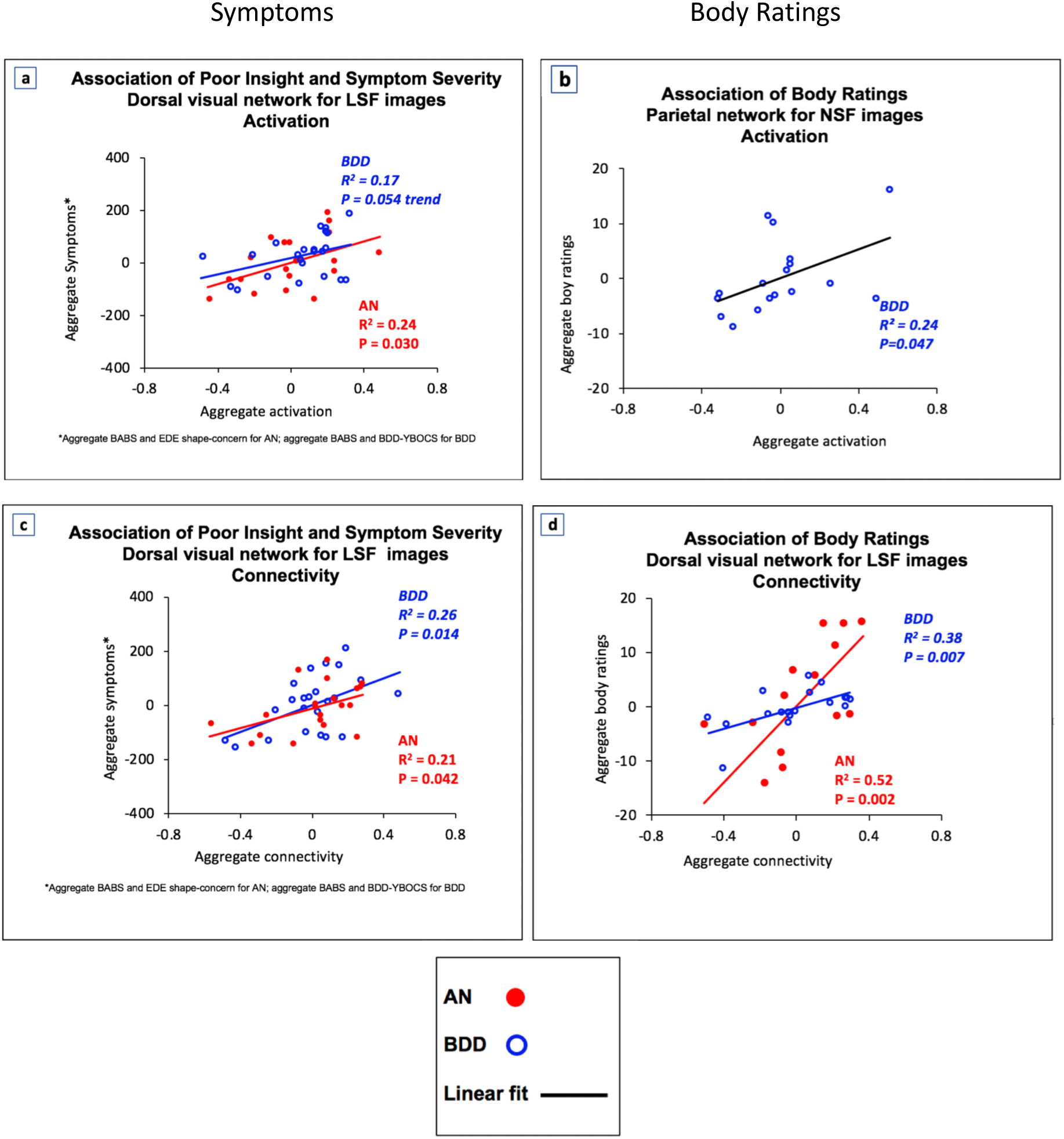
Brain activation and connectivity in dorsal visual and parietal networks are associated with poor insight, symptom severity, and body ratings^1^. For LSF images, dorsal visual network activation is associated with poor insight and symptom severity (4a, 4c). For NSF images, parietal network activation is associated with body ratings (4b). For LSF images, dorsal visual stream connectivity is associated with body ratings (4d). Note that the axes represent the values of latent space but do not relate linearly with the scales of the original variables. The fit lines represent a net aggregate directionality of the association of the latent independent and the latent dependent variables. Abbreviations: AN, anorexia nervosa; BABS, Brown Assessment of Beliefs Scale; BDD, body dysmorphic disorder; BDD-YBOCS, BDD Yale Brown Obsessive Compulsive Disorder Scale; EDE, Eating Disorder Examination. ^1^ Body ratings include attractiveness, aversiveness, and over/underweight judgments – see Supplemental Materials for details of ratings.

### Multivariate PLS associations of appearance ratings of bodies from brain activation

#### Dorsal Visual Network–LSF

For BDD, aggregate activation strength was associated with ratings of others’ bodies only at trend level (R^2^ =0.18, p=0.081).

#### Parietal Network–NSF (Fig. 4b)

For BDD, aggregate activation strength was associated with ratings of others’ bodies (R^2^ =0.24, p=0.047).

### Connectivity (Table 3)

#### Dorsal Visual Network–LSF

There was greater connectivity in both AN and BDD compared with controls in the right superior lateral occipital cortex (Figs. 5a, 5b). BDD showed greater connectivity than AN in the right lateral occipital cortex, right superior parietal lobule, right lingual gyrus and the right intracalcarine cortex (Fig. 5c). There were also areas of lower connectivity for BDD and AN compared with control in the right inferior frontal gyrus and left inferior frontal gyrus, respectively.

**Table 3.**
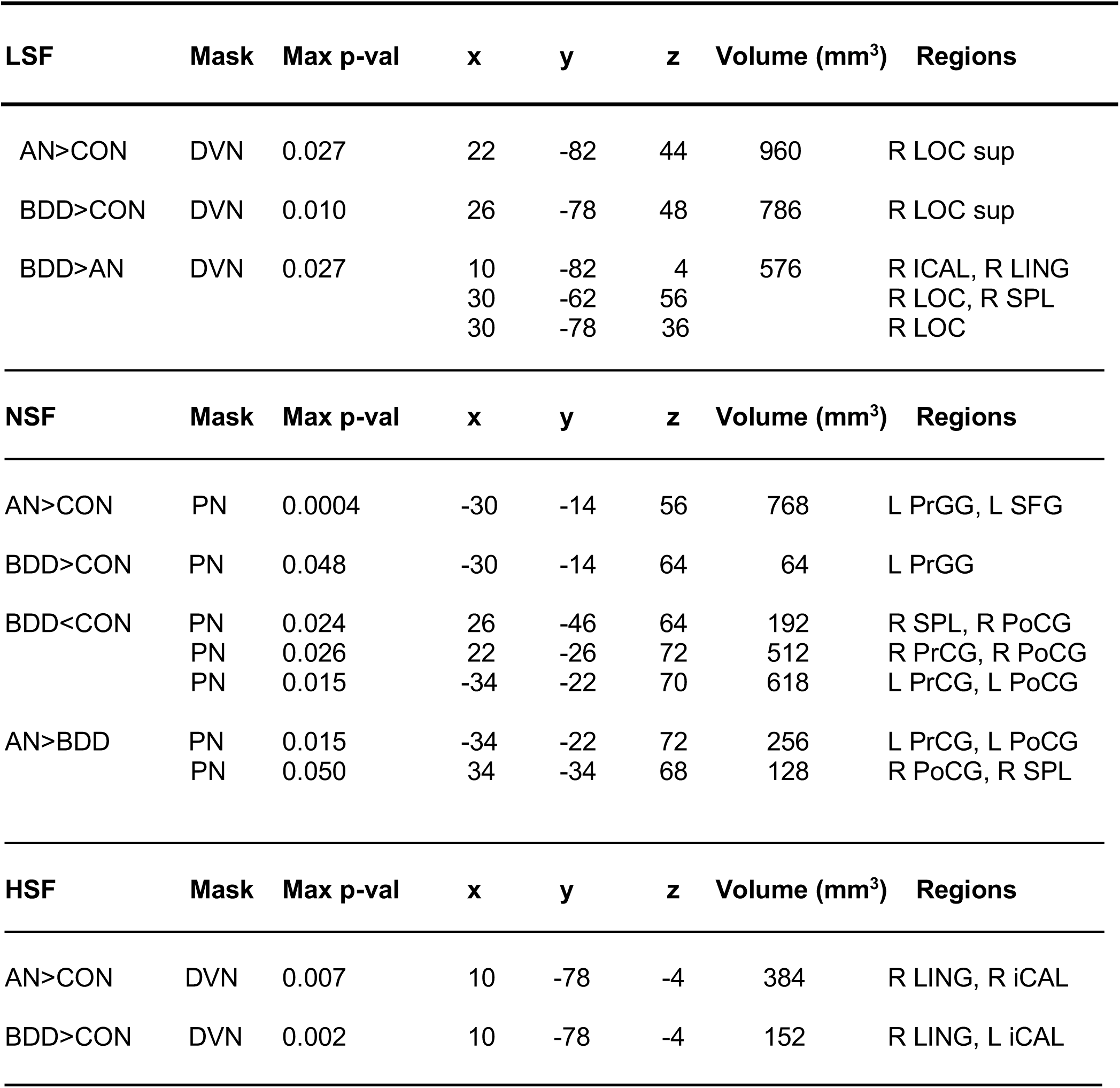
Coordinates of brain connectivity differences.

**Fig. 5.**
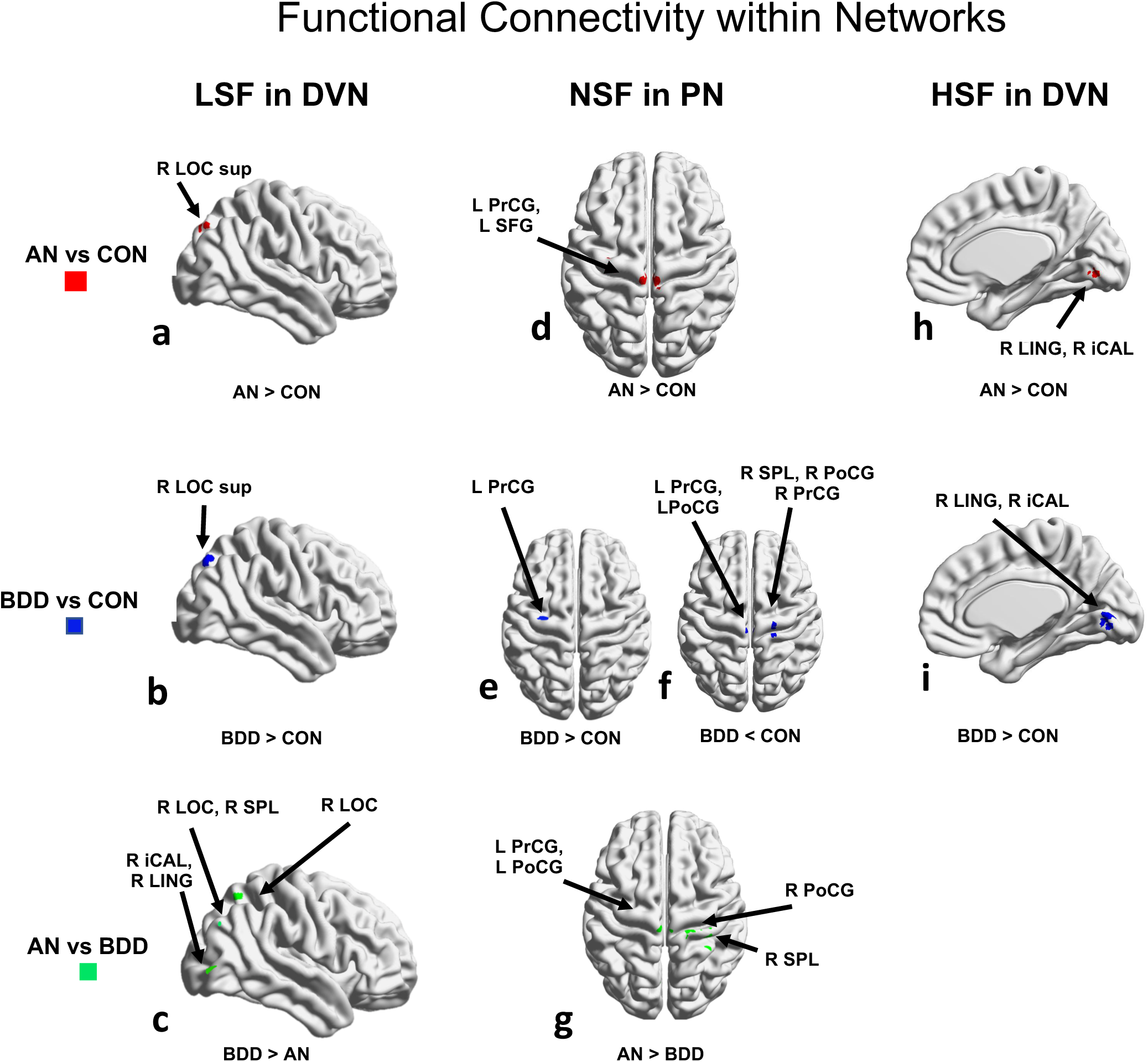
Functional connectivity differences. ANOVA identified regions differing among the 3 groups, using threshold-free cluster enhancement with a family-wise error correction at p<0.05, followed by pairwise group comparisons within significant regions. Between group results for low spatial frequency images in dorsal visual network are in 5a-c, for normal spatial frequency in parietal network are in 5d-g, and for high spatial frequency images in 4h-i. The rendered figures are registered to a 2mm MNI brain, but the analyses were performed in 4mm space. Abbreviations: AN, anorexia nervosa; BDD, body dysmorphic disorder; CON, controls; D, dorsal; F, frontal; L, left; R, right.

#### Striatal network-NSF

There were no significant differences between AN, BDD, and controls.

#### Parietal Network-NSF

Compared to control, AN and BDD had greater connectivity in the left precentral gyrus and left superior frontal gyrus (Figs 5d, 5e). BDD showed lower connectivity compared to controls in the right superior parietal lobule and pre- and postcentral gyri, bilaterally (Fig. 5f). AN showed greater connectivity than BDD in right superior parietal lobule and right postcentral gyrus (Fig. 5g).

### Exploratory HSF Connectivity

#### Dorsal Visual Network (Figs 5h, 5i)

Compared to control, AN and BDD had greater connectivity in the right lingual gyrus and right intracalcarine cortex. There were no significant differences between AN and BDD.

#### Parietal and Striatal networks

There were no significant differences between AN, BDD, and controls.

### Associations between symptom severity and brain connectivity

#### Dorsal Visual Network-LSF

Eating disorder severity (EDE total scores) in AN was inversely associated with connectivity in left occipital pole and left lateral occipital cortex (p=0.032, Fig. 3d). Follow-up exploratory analyses revealed that in AN shape concerns (EDE shape-concern subscale score) were inversely association with connectivity in left intracalcarine cortex (p=0.019, Fig. S3b).

### Multivariate PLS associations of symptom severity from brain connectivity

#### Dorsal Visual Network-LSF (Fig. 4c)

In BDD, aggregate connectivity was significantly associated with symptom severity (BABS and BDD-YBOCS scores; R^2^=0.26, p=0.014). In AN, there was also a significant relationship between connectivity and symptom severity (BABS and EDE shape concerns; R^2^=0.21, p=0.042).

### Multivariate PLS associations of appearance ratings of bodies from brain connectivity

#### Dorsal Visual Network-LSF (Fig. 4d)

In AN and in BDD, aggregate connectivity was significantly associated with ratings of others’ bodies (R^2^=0.52, p=0.002 for AN; R^2^=0.38, p=0.007 for BDD).

We repeated activity and connectivity PLS analyses after regressing out illness duration (AN and BDD) and lowest lifetime BMI (AN) (Supplemental Materials). The associations with the psychometrics did not change significantly.

## DISCUSSION

We compared and contrasted neural patterns of activation and connectivity for visual processing of bodies in two psychiatric disorders characterized by body image disturbance. Overall, in the dorsal visual and parietal networks, which are primarily involved in body visual processing, activation patterns in AN and BDD were largely different. BDD - but not AN - demonstrated hypoactivation compared with controls, and there were regions with both greater and lower activation in BDD compared to AN. However, there were more similarities in connectivity than activity in AN and BDD compared with controls. These similarities were in the dorsal visual network, with regions of primarily greater connectivity, and in the parietal network, with regions of both greater and lower connectivity. Contrary to our predictions, there were no significant activation or connectivity differences in the striatal network. Both activity and connectivity in the dorsal visual network were associated with clinical symptoms specific for AN and for BDD. Subjective ratings of appearance of others’ bodies were associated with activation and connectivity in the dorsal visual network in AN and BDD (trend level for activation for BDD) and in BDD with activity in the parietal network. This suggests that both AN and BDD may have links between neural functioning and perception that could involve aberrancies in visual sensory processing and (in BDD) higher order integrative systems.

### Aberrant visual system neural phenotypes

As hypothesized, there was significant hypoactivation in the dorsal visual network, specifically within the dorsal visual stream, for low-detail body images in the BDD group. Contrary to prediction, there was no significant hypoactivation in regions of this network in AN. The BDD result parallels previous findings of hypoactivity in BDD in visual systems for face perception (J. D. Feusner et al. 2010). The current results thus suggest similar deficiencies in processing global and configural aspects of low-detail bodies and low-detail faces. Interestingly, BDD also showed hypoactivation in dorsal visual stream regions compared to controls for NSF images (in portions of the parietal network that overlap with the dorsal visual stream) as well as for HSF images.

The absence of significant visual system activation abnormalities in AN is consistent with a previous study that found no significant differences from controls in visual systems when viewing others’ bodies (Sachdev et al. 2008). Several studies using own bodies, however, showed hypoactivation (reviewed in (Phillipou, Rossell, and Castle 2014; Suchan, Vocks, and Waldorf 2015)); yet, different patterns of findings could be attributed to others’ bodies being less personally salient than own bodies.

Contrary to predictions, AN and BDD demonstrated greater connectivity than controls in the dorsal visual network. This abnormal pattern was similar in both groups, particularly in later (superior lateral occipital cortex) parts of the visual processing stream, supporting a shared spatio-temporal phenotype involved in visual processing of bodies. We expected to find, as in previous connectivity studies for body processing, that AN would have lower connectivity in occipital and temporal regions that were associated with body-image distortions (Suchan et al. 2013). Another study found hyperconnectivity between precuneus and mid-temporal regions (Via et al. 2018). However, comparisons of these to the current study are challenging due to divergent methodologies.

In addition to these abnormalities in activation and connectivity compared to controls, there were important differences between persons with BDD compared to AN. BDD showed greater activation than AN in lingual gyrus (early visual processing stream) and greater connectivity in both early (intracalcarine and lingual gyrus) and later (superior lateral occipital cortex) portions of the dorsal visual network. In the parietal network BDD showed lower connectivity than AN in right superior parietal lobule and right postcentral gyrus. Although interpretation of these complex differences is not straightforward, lower activation and connectivity in visual systems in AN compared with BDD may indicate more consistent and severe disturbances for body perception in AN, consistent with bodies being more likely to have greater personal symptom relevance.

Both groups showed significant covariation between brain activity and clinical symptoms. Lower activation in early visual regions in AN were associated with worse eating disorder symptom severity, particularly worse body-shape concerns. Similar relationships were evident in AN for connectivity. Specifically, body-shape concerns were associated with connectivity in early visual systems while total eating disorder severity was associated with connectivity in occipital pole and lateral occipital cortex that represent later regions in the visual processing stream. In BDD, lower activation in dorsal visual network was associated with worse insight, while in AN the relationship was in the opposite direction. Moreover, in AN the association was in early visual systems, including the right lingual gyrus and intracalcarine cortex, while in BDD this association was observed in prefrontal systems, including the right inferior and middle frontal gyri (although outside of occipital and parietal cortices these regions were nevertheless connected with the dorsal visual network in our ICA masks).

As this was evident for LSF bodies, this pattern in AN may reflect that more severe impairment in holistic and configural processing at very early stages of visual processing would result in poorer ability to contextualize details of others’ bodies (particularly areas they preferentially attend to such as abdomen and thighs) relative to the whole body, and this is linked to their insight.

A possible explanation is that this pattern – if also operative when viewing their own body – may contribute to perceptual distortions for their own appearance. Those with the most impaired holistic and configural perception, particularly if occurring at early stages of visual processing, would be expected to have worse insight, because they do not have conscious control or awareness that their perception is distorted (J. Feusner, Deshpande, and Strober 2017).

In AN and BDD, multivariate analyses of regions of abnormal activation and connectivity in dorsal visual network showed associations with poor insight, core body dysmorphic symptoms for BDD, and eating disorder symptoms for AN. Therefore, aberrant neural function in multiple regions within the dorsal visual network may interact with each other to affect behavioral phenotypes, which themselves may interplay in manifold ways. The PLS results captured these complex relationships within potentially inter-related variables.

Further, similar relationships with connectivity were evident with ratings of pictures of others’ bodies’ attractiveness and aversiveness and how over/underweight they are. Both AN and BDD tend to rate others’ bodies as more overweight and less attractive than healthy controls (Moody et al. 2017). These results support the idea that aberrant visual system neural functioning could underlie disturbances in perception in both disorders.

### Abnormal activity and connectivity in the parietal network

In the parietal network, AN and BDD showed (predominantly) similar abnormal connectivity patterns; however, only BDD demonstrated abnormal activation patterns. There was lower activity in BDD in left superior parietal lobule and left superior lateral occipital cortex. Lateral occipital cortex was also hypoactive in the dorsal visual network, which may be due to overlapping spatio-temporal components making up dorsal visual and parietal networks. Lower activity within the parietal network could represent disturbances in parieto-occipital regions associated with detection, identification, and attention to body images, including one’s own (Hodzic et al. 2009) or another person’s body. The absence of significant activation differences between AN and controls within the parietal network when viewing *others’* bodies could reflect specificity of activation to *own*-body perception, but the exact mechanism for why our results differ from previous findings in other-body processing is unclear (Uher et al. 2005; Vocks, Busch, Gronemeyer, et al. 2010).

The regions of abnormal connectivity in AN and in BDD were primarily within somatosensory components of the parietal network. Previous connectivity studies in AN (Suchan et al. 2013; Via et al. 2018) probed other networks and regions, making comparisons with the current results difficult. However, evidence for distorted *own-body* representations in somatosensory cortex have been described in AN (Gadsby 2017). One speculation is that viewing others’ bodies might have triggered thoughts of self and thereby engage somatosensory components of the parietal network, previously demonstrated to occur in both AN and BDD (Moody et al. 2017).

Multivariate relationships between appearance evaluations and activity in the parietal network were evident in BDD, but not in AN. Higher-order integrative systems in the parietal network may impact perception of appearance in BDD in a way that is distinct from that in AN. Differences in activation and connectivity from the direct AN vs. BDD comparisons support this possibility. It remains unknown if these differences between AN and BDD (in activation, connectivity, and associations with clinical symptoms) reflect a qualitative difference in relationships between visual systems and perception, or if they reflect quantitative differences that could be due to differences in *own-body* appearance concerns across participants in our sample.

### Clinical implications

The partially overlapping clinical symptomatology and partially shared (although also distinct) disturbances in the dorsal visual network in AN and BDD have potential nosological and clinical implications. In BDD, hypoactivity within functionally connected regions comprising the dorsal visual network suggests reduced holistic integration and configural processing of spatial relationships of body images. This may underlie the tendency of those with BDD to “miss the big picture” when viewing appearance in general and to instead focus on specific aspects of body parts. Although significant hypoactivity was not evident in AN in the dorsal visual network, there were nevertheless significant univariate associations between lower activity in earlier systems within this network and worse eating disorder symptoms, which may have been driven specifically by shape concerns. The significant associations of connectivity (and activation for BDD) with appearance evaluations of others’ bodies also support the general clinical relevance of aberrant neural functioning being linked to analogous perceptual experiences and symptomatologies contributing to their respective body-image disturbances. Moreover, while categorical nosological systems including DSM-5 describe AN and BDD as separate disorders and include them in distinct superordinate categories, these results, in combination with previous results for viewing faces and houses (Moody et al. 2015; Li, Lai, Bohon, et al. 2015; Li, Lai, Loo, et al. 2015), suggest *partial* neurobiological similarities regarding visual processing.

Similar to some previous studies in AN that used others’ bodies (reviewed in (Phillipou, Rossell, and Castle 2014; Suchan, Vocks, and Waldorf 2015)), our findings support the premise that visual processing abnormalities are present when viewing others’ bodies. (This examination of others’ body processing is novel in BDD.) Therefore, perceptual abnormalities for bodies could be a general phenomenon rather than a phenomenon exclusive to one’s own appearance. The idea of pervasiveness of abnormalities is strengthened by our finding that activation and connectivity abnormalities were *disorder-relevant* but not *disorder-specific*, as opposed to self-body stimuli for AN and self-face stimuli for BDD.

Understanding the loci of pathological neural functioning that are associated with core symptoms of body image disturbance could ultimately contribute to improved and novel treatment methods (Beilharz et al. 2017; J. Feusner, Deshpande, and Strober 2017). While several studies have shown that body representation in AN can be malleable (Eshkevari et al. 2012; Keizer et al. 2014, 2016), body image distortion is a strong predictor for relapse in AN (Button 1986; Freeman et al. 1985). With further investigation, potential interventions in AN and BDD could include specific perceptual and behavioral retraining, as well as targeted brain stimulation, to remediate or compensate for abnormalities that may underlie perceptual distortions.

### Limitations & Future Directions

The AN group was weight-restored and analyses were limited to females in both groups; therefore, the generalizability to underweight AN and to males is unclear. Future studies would benefit from an increase in sample size, addition of underweight AN participants, and males. Our weight-restored AN participants were a mixture of those with either restricting or binge/purge subtypes. Future larger studies should examine possible differences between these subtypes. The duration of weight restoration data was not available for all AN participants to understand relationships to stages of recovery. The cross-sectional nature of this study prevents cause-and-effect to be established between aberrant neural functioning, clinical symptoms, and perceptual experiences. However, controlling for illness duration and lowest BMI did not appreciably change multivariate relationships between activity/connectivity and clinical symptoms or appearance ratings. Therefore, it is less likely that these findings resulted from a starvation state in AN, or from the effects of illness over time in either AN or BDD. Furthermore, several studies have shown that gray matter reduction from malnutrition in AN is reversed with weight restoration (Bernardoni et al. 2016; King et al. 2015; Seitz et al. 2014) so it is less likely that persistent starvation-related brain morphological effects could have influenced the functional results in the current study. Nevertheless, future longitudinal studies of the same cohort from the malnutrition state to weight-restoration are necessary for definitive answers about these questions. Finally, future research could address integration and segregation of networks (Ramsey 2018) to approach a more comprehensive and detailed understanding of body perception and processing.

## Conclusions

When viewing bodies, AN and BDD show distinct brain activation abnormalities and unique associations with disorder-specific symptoms. However, they share similar neural phenotypes of disrupted functional connectivity in the dorsal visual and parietal networks. These results in AN and BDD point to a complex pattern of aberrant activity and connectivity that is related to clinical symptoms and that may underlie analogous distorted perceptions of appearance.

## Acknowledgements

We would like to thank Amy Yu for contributions to the figures. Results were presented in part at the 56^th^ ACNP Annual Meeting, 2017.

## Compliance with Ethical Standards

### Funding

This study was funded by NIH: R01MH093535 (Feusner) and T32 GM08042 (Porterra/Kerr). The funding sources had no participation in conducting the study.

### Ethical approval

All procedures performed in studies involving human participants were in accordance with the ethical standards of the institutional and/or national research committee and with the 1964 Helsinki declaration and its later amendments or comparable ethical standards. The UCLA Institutional Review Board (IRB) approved the study.

### Informed consent

Informed consent was obtained from all individual participants included in the study.

