## SUPPLEMENTARY_MATERIALS for "Brain activation and connectivity in anorexia nervosa and body dysmorphic disorder when viewing bodies: relationships to clinical symptoms and perception of appearance"

**Brain Imaging and Behavior**

Teena D Moody, PhD[1]; Francesca Morfini, MS[2]; Gigi Cheng, BS[1]; Courtney L Sheen, MA[1]; Wesley Kerr, MD, PhD[3]; Michael Strober, PhD[1]; Jamie D Feusner, MD[1]

Affiliations:
[1]Department of Psychiatry and Biobehavioral Sciences, David Geffen School of Medicine at UCLA, Los Angeles, CA 90024-1759, USA, [2] Department of Psychology, Northeastern University, Boston, MA, USA, [3] Department of Neurology, David Geffen School of Medicine at UCLA, Los Angeles, CA 90095, USA

**Disclosures:** The authors report no competing interests. The funding sources had no participation in conducting the study. The UCLA Institutional Review Board (IRB) approved the study.

**Address for correspondence:**
Teena D Moody,,
https://orcid.org/0000-0001-7067-9512?lang=en
310-503-5630; UCLA Semel Institute 27-465, Los Angeles, CA 90095.

**Supplementary Methods**

1. **Description of metrics and illness duration**
2. **Inclusion and exclusion criteria**
3. **Body task design and subjective rating scales**
4. **Eigenvalues, coherence values and activation clusters**
5. **Mask creation**
6. **Dual regression analysis**
7. **PLS methods and parameters**
8. **MRI data preprocessing details**

**Supplementary Results:**

9. **Behavioral data from matching photos task in scanner**
10. **EDE shape-concern covariates**
11. **Lowest lifetime BMI and illness duration effects**

**Tables and Figures**

**Figure S1. Body ratings stimuli**

**Figure S2. Canonical networks masks**

**Figure S3. Associations with shape concerns for LSF images**

**Figure S4. Associations with symptom severity for HSF images**

**Table S1. Associations with symptom severity and insight scores**

**References for Supplementary Materials**

### Supplementary Methods

#### 1. Description of metrics and illness duration

BABS scores for 3 AN participants were imputed from age, MADRS, HAMA and EDE scores and used in our covariate and PLS analyses. Due to n=5 AN and n=5 BDD participants missing body rating scores, PLS analyses were performed on a subset of participants. We used age and age-squared as regressors of non-interest in all reported analyses to control for its effect, although results were substantially similar when age was not regressed out.

#### 2. Inclusion and exclusion criteria

We excluded any participants if they met the following criteria: 1) pregnancy; 2) ferromagnetic parts in their bodies; 3) heavier than 280 lbs.; 4) psychiatric medications within 8 weeks prior to enrolling in the study or currently undergoing cognitive behavioral therapy; 5) neurological disorder or any medical condition that may affect cerebral metabolism; 6) MADRS score >44; 7) BMI <18.5. Exclusion criteria for AN and BDD were: 1) current or past comorbidity with body dysmorphic disorder for AN, and with anorexia or bulimia nervosa for BDD; 2) any Axis I disorder other than dysthymia, major depressive disorder, panic disorder, agoraphobia, generalized anxiety disorder, or social phobia. These are highly common comorbidities in both disorders and it would not be a representative sample to exclude them. We excluded controls who had any Axis I diagnosis. Inclusion criteria for BDD and AN were: 1) meet criteria for BDD or AN; 2) BDD-YBOCS scores > 20 for BDD; 3) be weight restored (BMI>18.5) for AN.

#### 3. Body task design and subjective rating scales

All participants viewed digitized gray-scale frontal-view photographs of other people's bodies ranging from normal weight to overweight to provide a representation of average body types, for the matching task in the scanner for the fMRI experiment (Fig. 1 of manuscript). Following the fMRI experiment, outside of the scanner, participants rated the appearance of the same photos (Fig. S1). We used an equal sex ratio for the body stimuli to approximate the proportion of males and females in our society.

After the fMRI experiment, participants rated 16 bodies each of NSF, LSF, and HSF photographs, for a total of 48 bodies. There were two orders of randomly sorted bodies that were counterbalanced across participants. Participants were seated in front of a computer to view the images on the screen. There was no time limit on viewing and the investigator recorded their responses. Participants were asked to score bodies on measures of attractiveness and aversiveness on a scale of 0 to 10, with 0=very unattractive up to 10=very attractive and with 0=not aversive up to 10=extremely aversive. For NSF images, the mean attractiveness ratings for the unaltered images across groups was 5.0, 5.0, 4.7 for AN, BDD, CON, respectively ($F(2,49)=0.33$, $p=0.72$), and the mean aversiveness ratings were 3.5, 3.5, 2.2, ($F=3.48$, $p=0.04$). For LSF images, the mean attractiveness ratings for the unaltered images across groups was 4.6, 4.6, 5.0 for AN, BDD, CON, respectively ($F=(2,49)=0.39$, $p=0.68$), and the mean aversiveness ratings were 3.4, 3.9, 1.9, ($F=(2,49)=5.46$, $p=0.01$). Thus, on average, images were in a medium range between very unattractive and very attractive, and closer to not aversive than to extremely aversive. However, note that while AN and BDD participants did not differ from controls in attractiveness ratings, both groups found body images to be significantly more aversive than control participants for NSF and LSF images.

How overweight or underweight the body appears was judged on a scale from -10 for very underweight, 0 for normal weight, and 10 for overweight. Here the AN and BDD participants judged bodies as significantly more overweight than the control participants, for both NSF images, ($F(2,49)=4.83$, $p=0.012$) and LSF images, ($F(2,49)=7.14$, $p=0.002$) images. See Table 1 in manuscript for full results. Here we tabulated, but did perform analyses with the HSF subjective ratings. These participants are a subset of the participants reported in an earlier publication of body ratings (Moody et al. 2017).

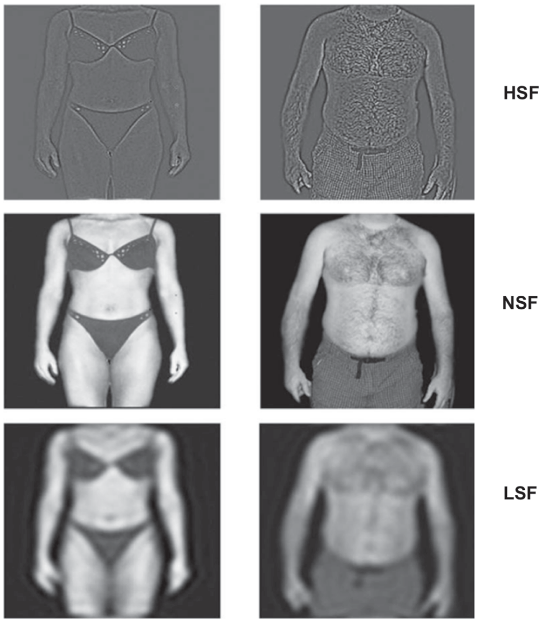


**Fig. S1.** Body ratings stimuli. The photos were non-copyrighted images from the internet.

### 4. Eigenvalues, coherence values and activation clusters

Eigenvalues represent the strength of the fMRI activation, which are mathematically calculated with their corresponding principal eigenvectors from a matrix of activation at every voxel for every time point within each regions of interest. Coherence values, derived from mean network connectivity scores for each subject, reflect the degree to which activation within specified brain regions covaries across time. We used eigenvalues and coherence values as metrics of brain activation brain and connectivity, respectively, for our PLS exploratory analysis.

For both AN and BDD we created 4 clusters from the LSF dorsal visual network (DVN) results for AN<CON and BDD<CON contrasts. Note that AN did not have clusters exceeding our GLM family-wise corrected results thresholded at $z>2.3$; therefore, for our exploratory analyses, we extracted clusters at $z>2.0$. We used FSL's cluster tool ([https://fsl.fmrib.ox.ac.uk/fsl/fslwiki/Cluster](https://fsl.fmrib.ox.ac.uk/fsl/fslwiki/Cluster)) to create clusters which we then binarized into masks to extract the eigenvalues from GLM or coherence values from dual regression results.

5. **Mask creation**

MELODIC was used to identify large-scale patterns of functional connectivity in all participants. Group-level independent component analysis (ICA) was performed to decompose our data set into 20 independent components, and to obtain network masks that were specific for NSF, LSF, and HSF bodies processing.

These masks were then correlated (Filippini et al. 2009) with canonical ICA networks (Smith et al. 2009) as described, Fig. S3. The components with the highest correlations for each spatial frequency with these canonical networks for DVN, PN, and SN were used in subsequent connectivity and activation analyses as our network-specific masks for analyses at each spatial frequency. All Pearson correlations values for selected network-specific independent components were greater than r=0.3. Canonical networks were Smith bm70 ic0032 and bm70 ic0036 for DVN, Smith bm10 ic0002 and bm70 ic0044 for the PN, and Smith bm20 ic004 for SN.

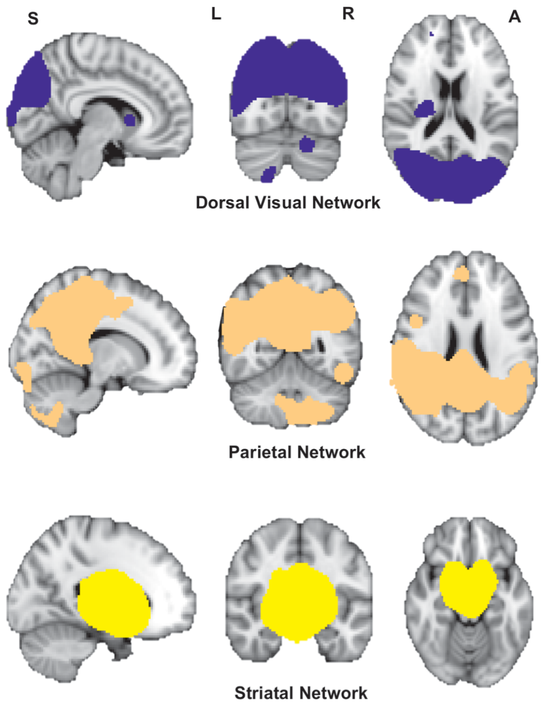


**Fig. S2.** Canonical ICA Network masks (Smith et al. 2009). Group-level independent components analysis (ICA) was performed to obtain network masks that were specific for bodies processing for each spatial frequency in our data set. Our masks were then correlated (Filippini et al. 2009) with these canonical networks.

6. **Dual regression analysis**

In this second stage (after the first stage of mask creation), we identified subject-specific temporal dynamics and associated spatial maps for each subject's fMRI data at LSF, NSF and HSF. This step used the full set of group-ICA spatial maps in a spatial regression for a linear fit against the separate fMRI data sets. The resulting time-course matrices comprising the temporal dynamics for each component and subject were then used in a temporal regression for a linear model fit against the fMRI data to estimate subject-specific spatial maps. These spatial map outputs of stage 2 were used to calculate coherence values for each subject within each network mask at each spatial frequency.

Finally, in stage 3, we calculated statistics using nonparametric permutation testing (FSL's Randomise function) using the 4-D component maps collected across subjects. Voxel-wise tests for statistically significant differences among groups used 5K-10K permutations (Winkler et al. 2014) depending upon critical values. The resulting spatial maps characterized the among-group and between-group differences controlling for age and age-squared. F-tests identified regions differing among the 3 groups, using threshold-free cluster enhancement with a family-wise error correction at $p<0.05$. Follow up between-groups comparisons were masked with the F-test results from the among-groups comparison.

7. **PLS methods and parameters**

PLS is a multivariate linear regression technique that projects variables into latent space components and then uses these components in a linear regression to determine associations with dependent variables. PLS can incorporate both direct and inverse relationships and represents the prediction as an aggregate effect. PLS is particularly suited when independent variables may be correlated, as can be the case for brain activation, connectivity, and/or psychometrics and ratings. (https://www.mathworks.com/help/stats/partial-least-squares.html***)***

For BDD, eigenvalues were extracted from 4 clusters centered in: right lateral occipital cortex, left postcentral gyrus/precuneous, left lateral occipital/left superior parietal lobule, and right precuneous. For AN, eigenvalues were extracted from 4 clusters centered in:

left lateral occipital cortex/angular gyrus, right lateral occipital cortex, left precuneous, and right middle temporal gyrus.

We used either eigenvalues or coherence values as independent/predictive variables in PLS analysis[1]. For the predictions of psychometrics scores, we used 4 independent variables (values from 4 DVN clusters) to predict 2 dependent variables, poor insight and symptom severity scores (BABS and EDE shape-concern for AN, BABS and BDD-YBOCS in BDD).

For the prediction of body ratings, we used the same 4 independent variables as above to predict 3 dependent variables (attractiveness, aversiveness, and under/overweight judgments); however, due to missingness in body ratings data, we used a reduced sample of n=16 (AN) and n=19 (BDD) for these analyses. Since eigenvalues and coherence values were extracted from results generated with age and age-squared (demeaned) as a confounder, we did not include age as a variable in any PLS models to avoid double corrections.

#### 8. MRI data preprocessing

Images were motion corrected, brain extracted, spatially smoothed using a Gaussian kernel of full-width at half-maximum of 5 mm, normalized, and high pass temporal filtered equivalent to 90 sec. Each functional image was registered to an MPRAGE, and then to Montreal Neurological Institute (MNI) standard brain space using FMRIB's Nonlinear Image Registration Tool. Image preprocessing for connectivity included resampling to 4mm space and motion-artifact correction by ICA-AROMA that denoised each individual participant's data by identifying and removing independent components attributed to noise as described elsewhere (Pruim et al. 2015) and recombining the remaining components into a single 4D denoised image.

### Supplementary Results

#### 9. Behavioral data from matching photos task in scanner

As expected, because the task is not difficult, mean accuracy across groups and across stimuli was high (95.7 ± 3.1% correct) and did not differ significantly among groups (repeated measures ANOVA: $F(2,57)=0.65$, $p=0.52$). There was a significant effect of stimulus type (spatial frequency) ($F(1,57)=4.98$; $p<0.03$), but the interaction between group and stimulus type was not significant ($F(2,57)=0.03$; $p=0.97$). Similarly, reaction times (mean=1.2 ± 0.3) did not differ among groups (repeated measures ANOVA: $F(2,57)=0.92$, $p=0.91$). There was a significant within-subject main effect of stimulus type ($F(1,57) = 6.96$; $p <0.01$), but no significant interactions between group and stimulus types ($F(2,57) = 0.74$; $p = 0.48$).

#### 10. EDE shape-concern covariates

Because shape and weight concerns are core aspects in AN and relevant for our behavioral task (Hartmann et al. 2015), we investigated the association of EDE shape-concern subscale scores in our activation and connectivity covariate analyses. We found that both activation and connectivity within DVN are inversely associated with EDE shape-concern (Figure S4). Shape-concern scores show a similar pattern of inverse activation as the total scores, but extending to bilateral intra- and supra-calcarine cortex, and occipital pole.

#### 11. Lowest lifetime BMI and illness duration effects

To mitigate the possibility that our results reflected effects of illness duration or severity of starvation state (lowest lifetime BMI), we regressed out these variables from eigenvalues and coherence values (illness duration and lowest lifetime BMI for AN, and illness duration for BDD) and obtained residualized values. Due to missing data we had n=16 AN and n=23 BDD with complete sets of values. To directly compare the impact of lowest BMI and illness duration per se, without confounding potential differences in the results due to an effect of the reduced sample size, we ran PLS on the reduced samples with raw data and with residualized values. The results were robust: the prediction of the psychometrics comparing these analyses did not significantly change, thus suggesting a minor impact of these variables on our results.

### Supplementary Figures and Tables

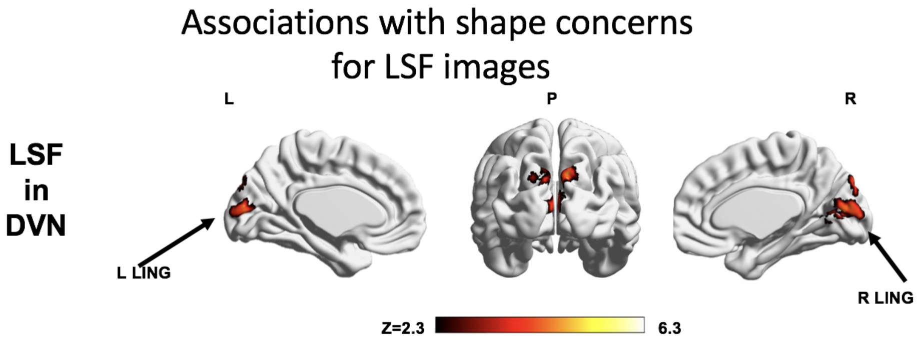


**a** **Inverse associations between EDE shape concern score and Brain Activation for Low Spatial Frequency Images**

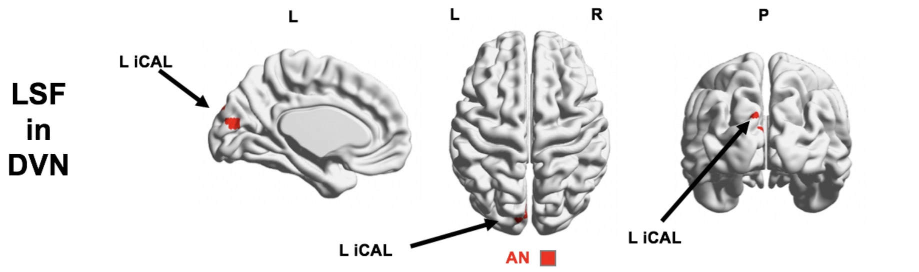


**b** **Inverse associations between EDE shape concern score *and Connectivity* for Low Spatial Frequency Images**

| **LSF - Activation** | **Z max** | **x** | **y** | **z** | **Regions** |
|---|---|---|---|---|---|
| EDE shape-concern (AN) | 3.96 | -20 | -84 | 6 | L LING |

| **LSF - Connectivity** | **Mask** | **p-val** | **x** | **y** | **z** | **Volume ($mm^3$)** | **Regions** |
|---|---|---|---|---|---|---|---|
| EDE shape-concern (AN) | DVN | 0.019 | 50 | 22 | 44 | 192 | L iCAL |

**Fig. S3.** Associations with shape concerns in the dorsal visual network for LSF bodies. a. Regions in which ***activation strength*** for LSF images inversely covary with EDE shape-concern scores. b. Regions in which ***connectivity strength*** for LSF images inversely covary with EDE shape-concern. Table shows coordinates of the activation and connectivity peak clusters for the within-group runs with EDE shape concerns as covariates of interest. Abbreviations: AN, anorexia nervosa; EDE, Eating Disorder Examination; iCAL = intracalcarine cortex; LING = lingual gyrus; R=right; L=left.

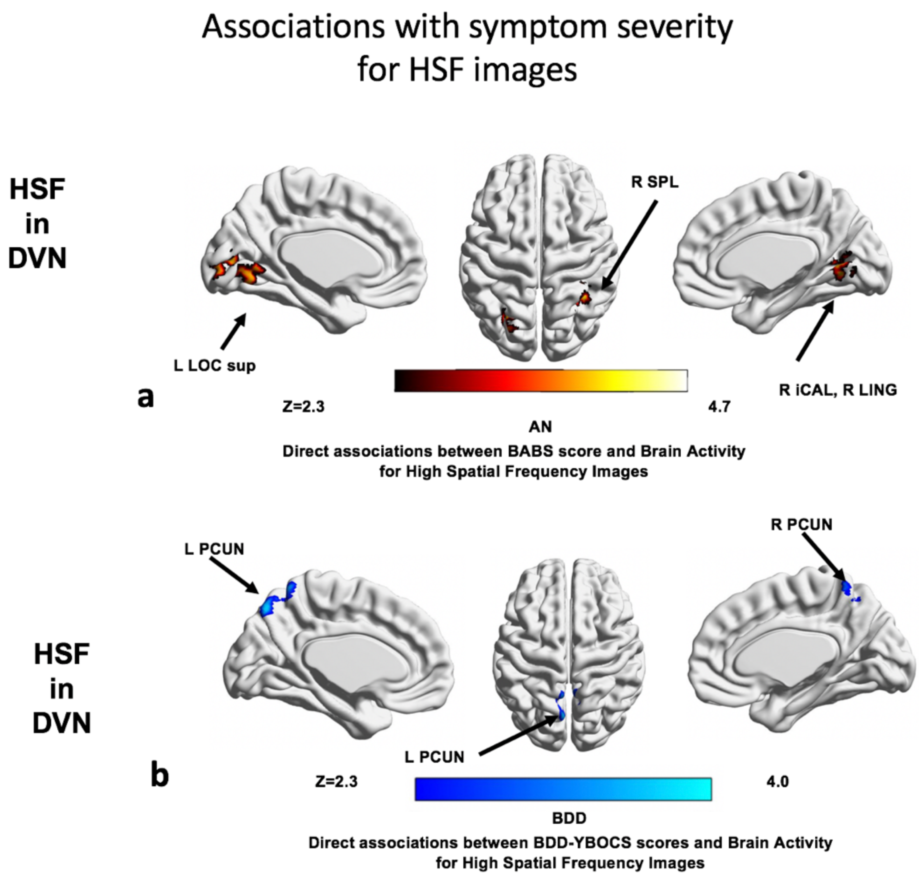


| HSF - Activation | Z max | x | y | z | Regions |
|---|---|---|---|---|---|
| BABS (AN) | 3.89 | 18 | -60 | 4 | R ICAL, R LING |
| (positive) | 4.57 | 32 | -42 | 50 | R SPL |
| | 4.02 | -28 | -64 | 38 | L LOC sup |
| BDD-YBOCS (BDD) | 3.98 | -2 | -72 | 52 | L PCUN |
| (positive) | 3.08 | 2 | -58 | 50 | R PCUN |

**Fig. S4.** Associations with symptom severity for HSF bodies stimuli.

Table shows the coordinates of the activation results for the within-group runs with psychometric covariates of interest. Table shows coordinates of the activation and connectivity peak clusters for the within-group runs with BABS and BDD-YBOCS as covariates of interest. Abbreviations: iCAL = intracalcarine cortex; LING = lingual gyrus; SPL=supraparietal lobule; LOC= lateral occipital cortex; PCUN=precuneus; sup=superior; R=right; L=left.

| LSF - Activation | Z max | x | y | z | Regions |
|---|---|---|---|---|---|
| BABS (negative) (BDD) | 4.02 | 54 | 16 | 32 | R IFG, R MFG |
| BABS (positive) (AN) | 3.52 | 14 | -64 | 2 | R LING, R iCAL |
| EDE total score (AN) | 3.30 | 0 | -86 | 8 | R&L SCAL, R&L iCAL |
| (negative) | 3.20 | 4 | -78 | 20 | R CUN, R SCAL |

| LSF – Connectivity | Mask | p-val | x | y | z | Volume ($mm^3$) | Regions |
|---|---|---|---|---|---|---|---|
| EDE total (AN) | DVN | 0.03 | -14 | -95 | 31 | 192 | L OCC, L OP |
| (negative) | | 0.05 | -46 | -81 | 12 | 128 | L LOC |

| NSF - Activation | Z max | x | y | z | Regions |
|---|---|---|---|---|---|
| BABS (negative) (BDD) | 3.58 | 2 | -38 | 30 | R PCC |

| HSF - Activation | Z max | x | y | z | Regions |
|---|---|---|---|---|---|
| BABS (positive) (AN) | 3.89 | 18 | -60 | 4 | R ICAL, R LING |
| | 4.57 | 32 | -42 | 50 | R SPL |
| | 4.02 | -28 | -64 | 38 | L LOC sup |
| BDD-YBOCS (BDD) | 3.98 | -2 | -72 | 52 | L PCUN |
| (positive) | 3.08 | 2 | -58 | 50 | R PCUN |

**Table S1. Symptom severity and insight results coordinates.** The table shows significance and coordinates of the general linear model and dual regression results for the within-group runs with psychometric covariates of interest. Positive = direct association, Negative = inverse association. Abbreviations: R = right; L= left; AN, anorexia nervosa; BABS, Brown Assessment of Beliefs Scale; BDD, body dysmorphic disorder; EDE, Eating Disorder Examination; Z, z scores; ANG = angular gyrus; iCAL = intracalcarine cortex; IFG = inferior frontal gyrus; LING = lingual gyrus; LOC= lateral occipital cortex; MFG = middle frontal gyrus; OP = occipital pole; PAC = paracingulate gyrus; PrCG = precentral gyrus; PCC = posterior cingulate gyrus; PCUN = precuneous; SPL= supraparietal lobule; SCAL = supracalcarine cortex; sup = superior division.
